# *In vivo* imaging of radial spoke proteins reveals independent assembly and turnover of the spoke head and stalk

**DOI:** 10.1101/240143

**Authors:** Karl F. Lechtreck, Ilaria Mengoni, Batare Okivie, Kiersten B. Hilderhoff

**Affiliations:** Department of Cellular Biology, University of Georgia, Athens 30602, GA

**Keywords:** axoneme, cilia, flagella, intraflagellar transport (IFT), RSP3, RSP4

## Abstract

Radial spokes (RSs) are multiprotein complexes regulating dynein activity. In the cell body and ciliary matrix, RS proteins are present in a 12S precursor, which is converted into axonemal 20S spokes consisting of a head and stalk. To study RS assembly *in vivo*, we expressed fluorescent protein (FP)-tagged versions of the head protein RSP4 and the stalk protein RSP3 to rescue the corresponding *Chlamydomonas* mutants *pfl*, lacking spoke heads, and *pf14*, lacking RSs entirely. RSP3 and RSP4 mostly co-migrated by intraflagellar transport (IFT). Transport was elevated during ciliary assembly. IFT of RSP4-FP depended on RSP3. To study RS assembly independently of ciliogenesis, strains expressing FP-tagged RS proteins were mated to untagged cells with, without, or with partial RSs. RSP4-FP is added a tip-to-base fashion to preexisting *pf1* spoke stalks while de novo RS assembly occurred lengthwise. In wild-type cilia, the exchange rate of head protein RSP4 exceeded that of the stalk protein RSP3 suggesting increased turnover of spoke heads. The data indicate that RSP3 and RSP4 while transported together separate inside cilia during RS repair and maintenance. The 12S RS precursor encompassing both proteins could represent transport form of the RS ensuring stoichiometric delivery by IFT. (196 of 200)

## Introduction

The 9+2 axoneme is a highly ordered molecular machine conferring motility to cilia and flagella. While providing stability to several μm-long flagella of just 200-nm in width, the axoneme is flexible to execute complex waveforms with ~100 hz. Mutant analysis and proteomic studies have identified a broad repertoire of axonemal proteins, which are synthesized in the cell body and transported into the cilia (Lefebvre and Silflow, 1999; Pazour et al., 2005). How the cell body precursors of the axoneme are transported and assembled remains largely unknown. Here, we study the assembly of radial spokes (RSs), which project from the nine doublet microtubules toward the central pair and regulate the activity of dynein by transmitting mechanical cues from the central pair (Oda et al., 2014). Mutations affecting the RSs result in paralyzed flagella (pf) phenotypes in protists (Piperno et al., 1977; Vasudevan et al., 2015) and cause primary ciliary dyskinesia in humans (Frommer et al., 2015). The RS consists of ~23 distinct proteins with diverse functional domain including several protein-protein interaction and signaling protein motifs (Luck et al., 1977; Yang et al., 2006). During ciliary assembly, RS precursors move from the cell body into the growing organelle. Intraflagellar transport (IFT), a bidirectional motor-based motility of protein carriers has been implicated in RS transport but direct evidence is still missing (Johnson and Rosenbaum, 1992; Qin et al., 2004).

Certain axonemal substructures move into cilia in form of multiprotein complexes. Inner and outer dynein arms, for example, are preassembled in the cell body into complexes thought to contain all subunits present in the corresponding mature structures inside cilia (Fowkes and Mitchell, 1998; Viswanadha et al., 2014). Outer dyneins arms (ODAs) isolated from the cell body will dock at the right sites onto the axoneme *in vitro* indicating that they are assembly competent and probably functional. In contrast, RS assembly consists of two phases: First, a 12S complex is formed in the cell body and transported into cilia (Qin et al., 2004). Inside cilia, the 12S complex is converted into the mature 20S complex and docked onto the axonemal microtubules (Diener et al., 2011). During spoke maturation, additional RS proteins that are not part of the 12S precursor (e.g., RSP16, HSP40 and LC8) are incorporated (Gupta et al., 2012). However, how the Γ-shaped 12S particles is converted into the T-shaped 20S RSs is unclear. The T-shaped RSs consist of the spoke head, the spoke neck, and the spoke stalk with at least six, three, and nine subunits, respectively. A key structural protein of the stalk is RSP3, whose N-terminus is close to microtubule-associated base of the spoke while the C-terminus extends into the spoke head (Oda et al., 2014). Mutants lacking RSP3 assemble spoke-less axonemes. The two wings of the spoke head are formed by homodimers of the paralogous proteins RSP4 and RSP6, respectively. Mutants in RSP4 lack the spoke heads while the stalks are retained. Both head and stalk proteins are part of the 12S RS precursor. How the 12S complex is converted into the mature 20s RS remains unclear.

Here, we used fluorescent proteins (FP)-tagged RS proteins in *Chlamydomonas reinhardtii* to analyze the transport and assembly of the RS. The head protein RSP4 and the stalk protein RSP3 are transported by IFT and both proteins mostly co-migrate. However, both proteins are added with distinct patters to the axoneme during RS assembly, turnover and repair. Our data indicate that RSP3 and RSP4 jointly enter the cilia, presumably in the previously characterized 12S complex, but separate inside the cilium during RS repair and turnover. We propose that the 12S RS precursor presents a transport complex of RS proteins.

## RESULTS

### RSP4-sfGFP rescues the spoke-less pf1 mutant

Using a derivate of the bicistronic vector pBR25, the spoke-head protein RSP4 was tagged at its C-terminus with sfGFP and expressed in *C. reinhardtii pf1*, a mutant lacking endogenous RSP4 (Fig. 1A; (Rasala et al., 2013)). As previously reported for RSP4-GFP, expression of RSP4-sfGFP rescued the paralyzed flagella (*pf*)-phenotype in ~50% of the transformants analyzed (Fig. 1B; (Oda et al., 2014)). TIRF imaging of live cells showed that RSP4-sfGFP was present along the entire length of the flagella with exception of the very tip (Fig. 1C). For western blot analyses, we used an antibody raised against RSP6, which also reacts with RSP4. RSP4 and RSP6 are paralogues sharing 47% identity and depend on each other for assembly into flagella (Curry et al., 1992). In cilia of control cells, the antibody recognized two bands corresponding to RSP4 and the somewhat smaller RSP6 (Fig. 1D). Both bands were absent in the *pf1* mutant cilia. In the *pf1* RSP4-sfGFP rescue strain, the RSP6 band was retained while the RSP4 band was replaced by two slower migrating bands. These bands also reacted with anti-GFP and hence represent RSP4-sfGFP and uncleaved ble-RSP4-sfGFP; the amount of uncleaved ble-RSP4-sfGFP varied between strains and preparations (Fig. 1b, d). In summary, expression of RSP4-sfGFP restored wild-type motility in *pf1* suggesting that it can be used to monitor radial spoke (RS) transport in living cells.

**Figure 1).**
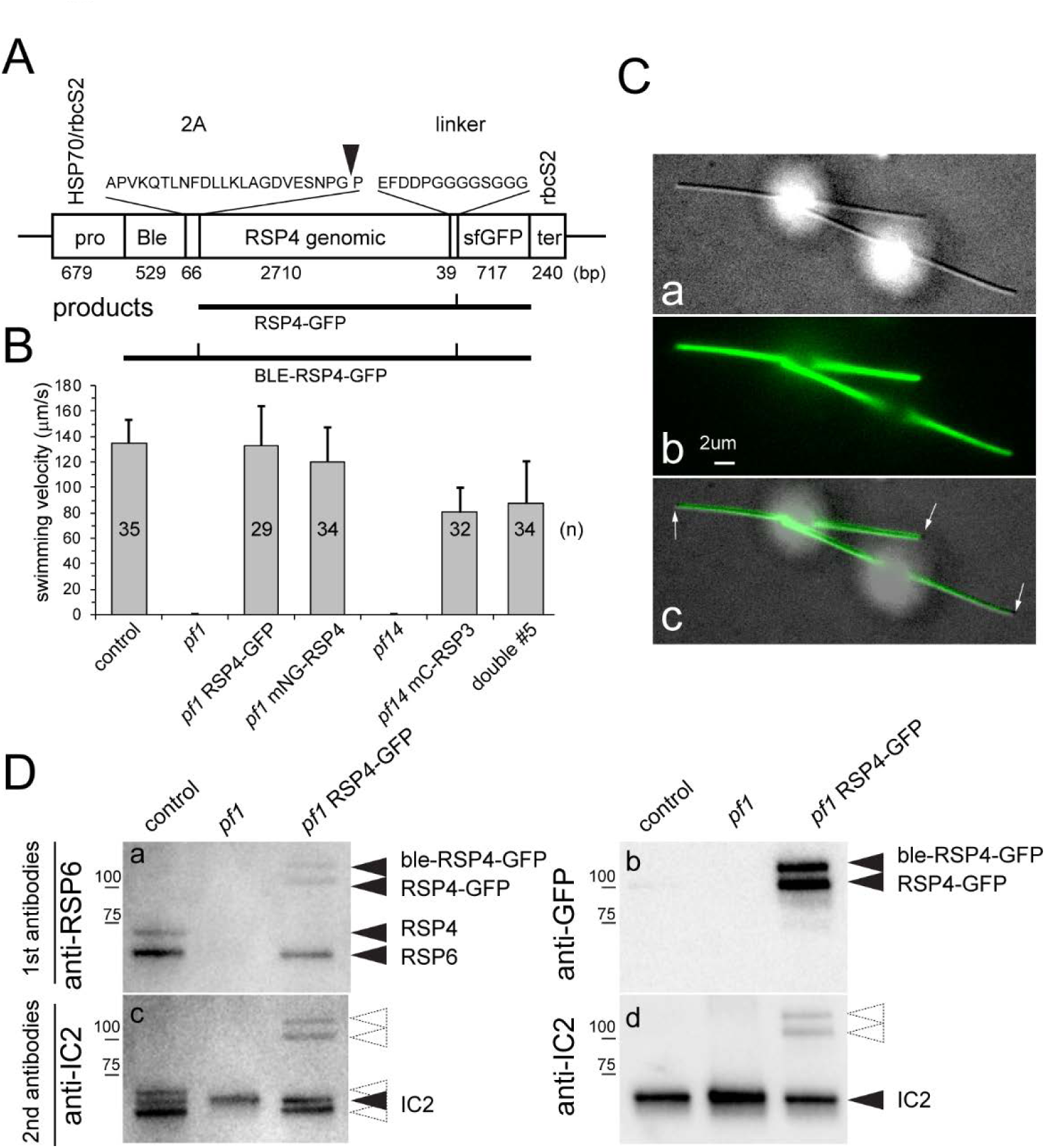
RSP4-sfGFP rescues the pf14 mutant. A) Schematic presentation of the bicistronic vector used to express RSP4-sfGFP. Due to incomplete cleavage of the 2A sequence, the transformants expressed both RSP4-sfGFP and uncleaved ble-RSP4-sfGFP. B) Swimming velocity of wild type, and the *pf1* and *pf14* rescue strains. n, number of cells analyzed. Note incomplete rescue of the motility phenotype in strains expressing mC-RSP3. C) DIC (a), TIRF (b), and merged (c) image of a live cell expressing RSP4-sfGFP. Arrows indicate the fkafellar tips lacking RSP4-sfGFP signal. Bar = 2μm. D) Western blots analyzing flagella isolated from wild-type, *pf1*, and the *pf1* RSP4-sfGFP rescue strain. The two replicate membranes (a/c and b/d) were first stained with anti-RSP6 (a) or anti-GFP (b) and subsequently with anti-IC2, a component of outer arm dynein, as a loading control. The positions of RSP6, RSP4, RSP4-sfGFP, ble-RSP4-sfGFP, and standard proteins are indicated. Dashed arrowheads indicate residual signals from the first round of antibody staining.

### RSP4-sfGFP is a cargo of IFT

Previous data indicate that radial spokes move via IFT into cilia but the direct observation of this transport has not yet been reported (Johnson and Rosenbaum, 1992; Qin et al., 2004). After photobleaching of RSP4-sfGFP already incorporated into the flagella, IFT-like transport of RSP4-sfGFP became visible (Fig. 2A). In steady-state flagella, anterograde IFT of RSP4-sfGFP was observed only occasionally with a frequency of 0.6 transports/min. To study transport in growing flagella, cells were deflagellated by a pH shock and allowed to initiate flagellar regeneration prior to mounting for TIRF imaging. In such cells, the average anterograde transport frequency of RSP4-sfGFP was 10.4 transports/min and a maximum of 30 transports/min was observed (Fig. 2B, C). We conclude that RSP4-sfGFP is a cargo of IFT and that RSP4 transport is strongly upregulated in regenerating flagella.

**Figure 2).**
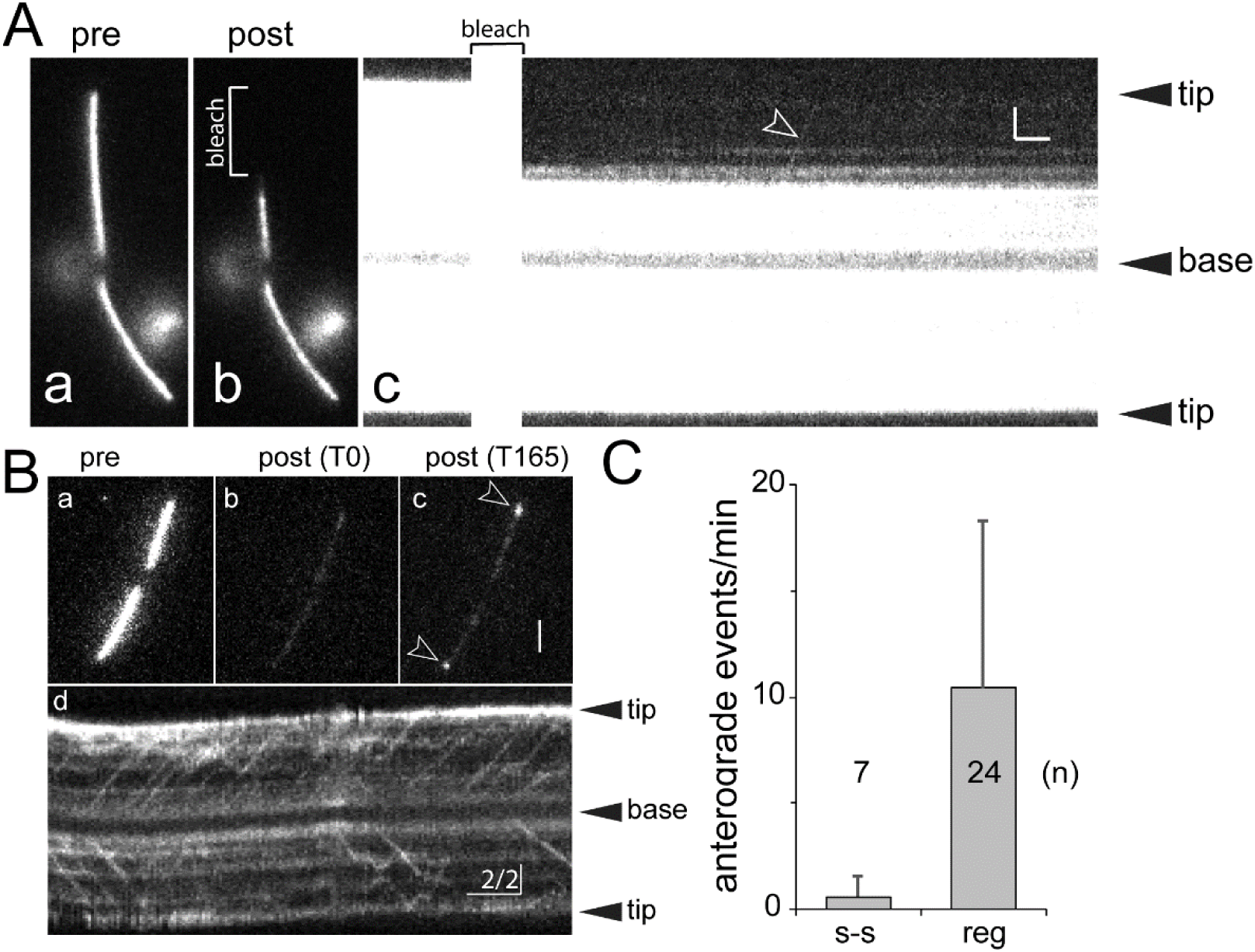
Transport o/RSP4-s/GFP is upregulated during flagellar growth. A) Cell with steady-state flagella before (a) and after (b) photobleaching of the distal segment of one flagellum (indicated by a bracket in b). The corresponding kymogram shows a single anterograde transport event (arrowhead in c). Bars = 2s 2μm. B) Cells during flagellar regeneration before (a), immediately after photobleaching (b, T0), and 165 s after the bleach (c). Note incorporation of unbleached RSP4-sfGFP at the distal ends of both flagella (arrowheads in c) indicative for axonemal elongation and spoke assembly. d) Kymogram of a regenerating flagellum showing numerous transport events of RSP4-sfGFP. Bars = 2s 2μm. C) Anterograde IFT events of RSP4-sfGFP observed in steady-state (s-s) and regenerating (reg) flagella. n, number of flagella analyzed.

### *RSP4 incorporates with distinct patterns into wild-type*, pf1, *and* pf14 *flagella*

In dikaryons obtained by mating *pf14* and wild-type gametes, the repair of RS-deficient *pf14* flagella progresses from the tip toward the base as assessed using antibodies to RSP3 and RSP2 (Johnson and Rosenbaum, 1992). The experiments suggest that radial spokes are transported by IFT to the flagellar tip, where they are released followed by diffusion to axonemal docking sites. To study radial spoke assembly *in vivo*, the *pf1* RSP4-sfGFP strain was mated to wild-type, *pf1*, and *pf14* gametes (Fig. 3A). The zygotes possessed two GFP-positive flagella derived from the *pf1* RSP4-sfGFP donor strain and two acceptor flagella initially devoid of GFP-tagged protein. In *pf1* RSP4-sfGFP x *pf1* zygotes, RSP4-sfGFP was added in a pronounced tip-to-base pattern to the formerly *pf1* flagella (Fig. 3A a-b). Most cells showed RSP4-sfGFP incorporation also in a small region near the flagellar base, a region for which IFT-independent assembly of RSs has been proposed (Alford et al., 2013). Older zygotes are recognized by the presence of shorter flagella, which progressively shorten during zygote maturation (Lechtreck et al., 2013). In such zygotes, RSP4-sfGFP was equally or almost equally present along all four flagella indicative for complete repair. The addition of RSP4-sfGFP to pf14-derived flagella of *pf1* RSP4-sfGFP x *pf14* zygotes follows a distinct pattern: While often stronger near the tip, RSP4-sfGFP incorporation was also present along the length of the pf14-derived flagella (Fig. 3A d-f). In matings with wild type, RSP4-sfGFP was present in a spotted fashion along wild-type derived flagella (Fig. 3A g-j). Kymograms revealed that these signals are stationary indicating stable docking of the protein to the axoneme (Fig. 3B). More continuous signals representing RSP4-sfGFP were detected in the wild-type derived cilia of older zygotes (Fig. 3A j). We conclude that untagged RSP4 was exchanged with RSP4-sfGFP indicative of spoke head turnover. With ~2 events/min, the transport frequency of RSP4-sfGFP in zygotic flagella was above that of non-regenerating flagella and below that of regenerating flagella. Regardless of whether RSs were present, partially present (p/1), or absent (*pf14*), the transport frequencies were similar indicating that RSP4-sfGFP transport is not regulated by the presence or absence of intact RSs inside cilia (Fig. 3C).

**Figure 3).**
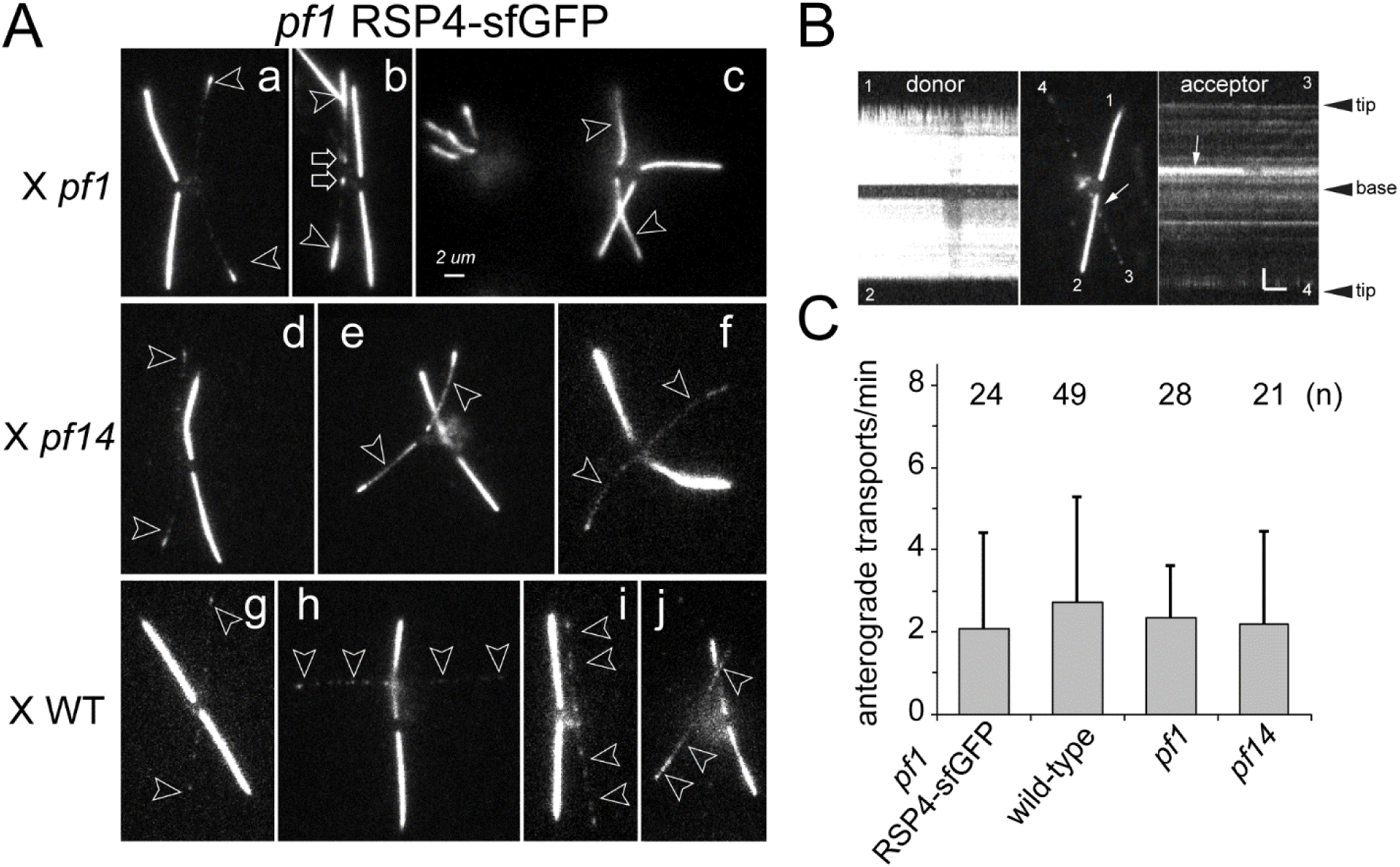
RSP4-sfGFP incorporates with distinct patterns into pf1 and pf14 flagella. A) Gallery of still images of life zygotes from crosses between *pf1* RSP4-sfGFP and *pf1* (a-c), *pf14* (d-f), and wild type (WT, g-j). Acceptor flagella are marked by arrowheads. Late zygotes are shown in c; arrows in b mark proximal RSP4-sfGFP signals. Bar = 2μm. B) Kymograms and still image of a *pf1* RSP4-sfGFP x wild type zygote. Horizontal trajectories in the acceptor flagella reveal that RSP4-sfGFP is stationary likely because of docking to the axoneme. Arrow: Crossing point of flagellum No. 2 and No. 3. Bar = 2 s 2 μm. C) Transport frequency of RSP4-sfGFP in zygotic flagella. n: number of flagella analyzed.

### RSP4-sfGFP transport is highly processive

The acceptor flagella of zygotes are well suited to image details of RSP4-sfGFP transport: They are full-length allowing to study RS assembly independently of ciliogenesis and initially lack RSP4-sfGFP incorporated in the axoneme making photobleaching unnecessary (Fig. 4A). Most RSP4-sfGFP transport events progressed uninterrupted from the flagellar base to the tip (Fig. 4A, C). Anterograde transport of RSP4-sfGFP progressed at typical IFT rates with an average velocity of 2.4 μm/s (STD 0.4 μm/s, n=256); retrograde transport was observed only rarely (Fig. 4A, B). Transitions from IFT to diffusion and vice versa along the flagellar shaft were observed as well (Fig. 4A b). RSP4-sfGFP arriving via IFT at the flagellar tip often remained stationary for extended times (Fig. S1). In detail, RSP4-sfGFP remained in a distal position for ~1s (STD 0.4s, n=9) after arriving via IFT at the tip before moving to a subdistal position, in which it remained for ~20 s or more (Fig. 4A; recordings typically lasted for 30–60s). Such tip-bound particles were observed to regaining movement (Fig. 4A d) suggesting RSP4-sfGFP often dwells for extended periods near the tip before moving into the flagellar shaft. Apparent docking of RSP4-sfGFP onto *pf1* axonemes was rarely observed; the putative docking events shown in Fig. 4A were preceded by extended periods of diffusion (Fig. 4A d). To summarize, RSP4-sfGFP transport during the repair of *pf1* flagella is characterized by processive IFT to the tip, prolonged dwelling at the tip, and extended diffusion prior to docking along the axoneme.

**Figure 4).**
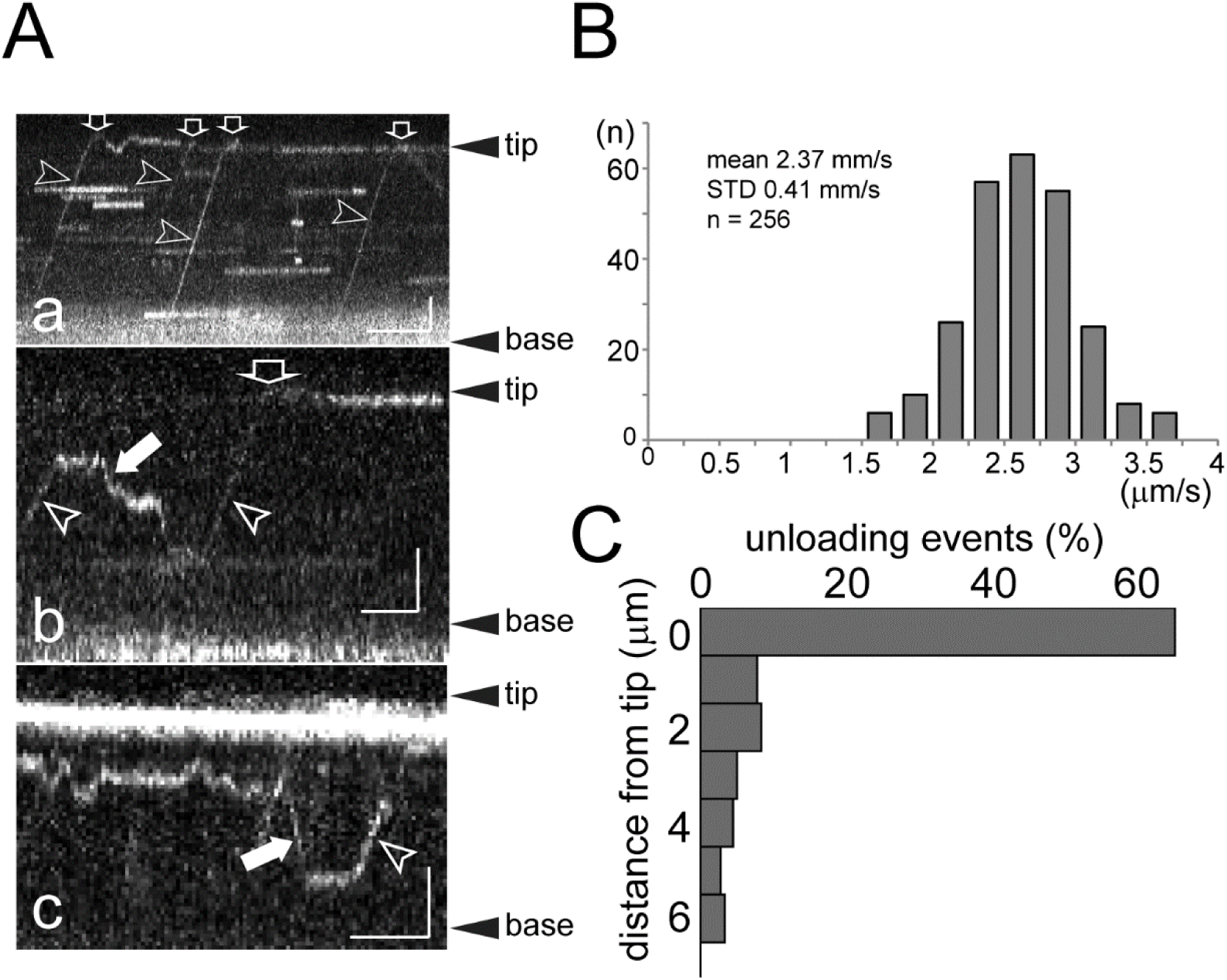
RSP4-sfGFP is mostly transported to the flagellar tip. A) Gallery of kymograms depicting IFT transport to the flagellar tip (a), unloading and reloading (b), dwelling at the tip (c), and retrograde transport. Arrowheads, anterograde transports; filled arrows, retrograde transport; open arrows, arrival at the ciliary tip. Bars = 2s 2μm. See figure for additional examples. B) Distribution of the velocity of anterograde RSP4-sfGFP transports. C) Distribution of IFT transport termination points of RSP4-sfGFP along the flagella. Note high share of RSP4-sfGFP transport moving processively to the flagellar tip.

### RSP3 is a cargo of IFT

Next, we rescued the *pf14* mutant by expression of RSP3 tagged at its N-terminus with mNeonGreen (mNG); rescue was also observed for mScarlet-I (mS), mCherry (mC), and mTAG-BFP fusions (Fig. 5A). Similar to RSP4-sfGFP, mNG-RSP3 is transported by IFT; its transport frequency is elevated during flagellar regeneration, and most transports progressed uninterrupted to the flagellar tip (Fig. 5B). At the flagellar tip, mNG-RSP3 remained briefly stationary before the on-set of diffusion. In contrast to RSP4-sfGFP, an extended stationary phase near the tip was not observed for mNG-RSP3 (Fig. 5B). In matings with *pf14*, mNG-RSP3 was incorporated along the length of the *pf14*-derived flagella; incorporation was somewhat stronger in the distal flagellar region (Fig. S2).

**Figure 5).**
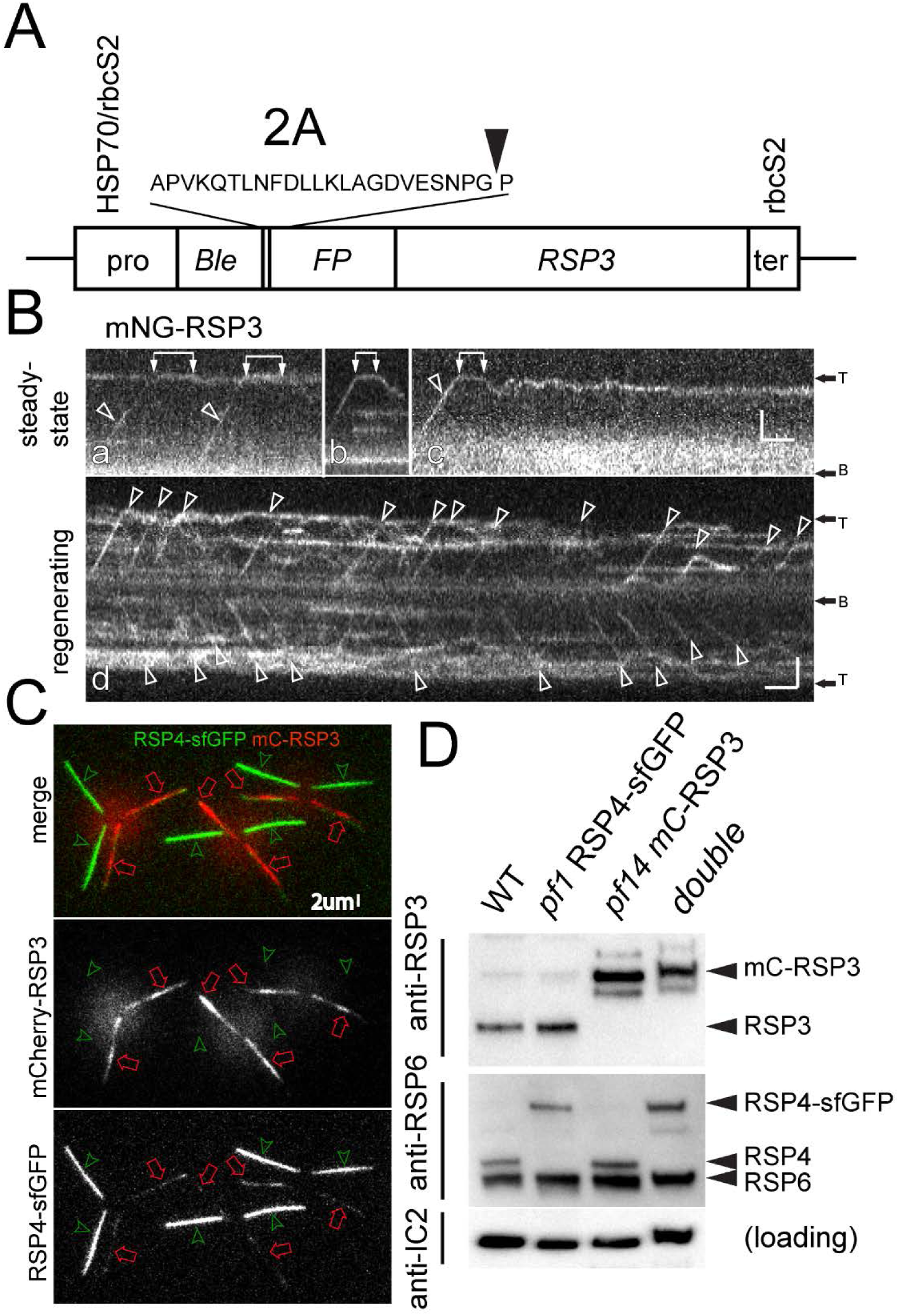
Generation o/strain with two-color radial spokes. A) Schematic presentation of the vector used for FP-tagging of RSP3. B) Kymograms showing mNG-RSP3 in steady-state (a-c) and regenerating (d) flagella. Transport events are marked with arrowheads, interconnected pairs of white arrows indicate dwelling of mNG-RSP3 at the flagellar tip. Bars = 2s 2μm. C) Still image of three life *pf1* RSP4-sfGFP x *pf14* mC-RSP3zygotes. *pf1* RSP3-sfGFP and *pf14* mC-RSP3 derived flagella are marked by green arrowheads and red arrows, respectively. Note incorporation of RSP4-sfGFP along the length of the pf14 mC-RSP3 derived flagella. Bar = 2μm. D) Western blots analyzing flagella isolated from wild-type (WT), *pf1* RSP4-sfGFP, *pf14* mC-RSP3, and *pf1 pf14* RSP4-sfGFP mC-RSP3 (double) cells. Two replica membranes were stained with anti-RSP3 and anti-RSP6 as indicated. Anti-IC2 staining was used to control for equal loading.

Biochemical studies in *C. reinhardtii* showed that RSP4 and RSP3 are both present in the 12S RS precursor complex in the cell body (Diener et al., 2011; Qin et al., 2004). To analyze whether these proteins remain associated during transport and assembly, we generated a strain expressing RSP3 and RSP4 tagged with different FPs by mating *pf14* mC-RSP3 to *pf1* RSP4-sfGFP. The zygotes initially possessed two red and two green flagella (Fig. 5C). At later time points, RSP4-sfGFP was readily detected in the *pf14* mC-RSP3-derived flagella whereas only traces of mC-RSP3 were observed in the *pf1* RSP4-sfGFP derived flagella (Fig. 5C). The observation suggests that RSP3 and RSP4 in the wild-type acceptor flagella are exchanged at distinct rates with the tagged proteins. A strain expressing both RSP4-sfGFP mC-RSP3 in the corresponding double mutant background was identified among the progeny of this cross by TIRF microscopy and western blotting (Fig. 5D).

### During the repair of pf1 axonemes, RSP4 complexes are added to preexisting stalks

Flagella of *pf1* already possess RSP3-containing spoke stalks. The repair of *pf1* axonemes in dikaryons could occur by the addition of spoke heads onto the preexisting stalks or by replacement of those stalks with newly imported entire RSs; the outcome will potentially inform on the stability of the RSP3/RSP4-containing RS precursor (Fig. 6A). To distinguish between these two possibilities, the mC-RSP3 RSP4-sfGFP double strain was mated to *pf1* and wild-type gametes. In wild type-derived flagella of such zygotes, both RS proteins were visible near the flagellar tip indicative for a limited turnover of the distal axonemal segment as previously reported for tubulin (Fig. 6B a, b; (Harris et al., 2016; Marshall and Rosenbaum, 2001). As described above, RSP4-sfGFP was also present in a spotted fashion along the length of wild type-derived flagella indicative for the replacement of untagged RSP4 with the tagged protein. In *pf1*-derived flagella, RSP4-sfGFP was readily visible in distal segments of early zygotes and along most of the length in later zygotes indicative for tip-to-base assembly (Fig. 6B c-e). In contrast, the signals representing mC-RSP3 in the mutant-derived flagella were generally weak including those of late zygotes, which contained a near full complement of RSP4-sfGFP (Fig. 6B e). Apparently, RSP4-sfGFP can be added to *pf1* flagella independently of mC-RSP3 incorporation. The data favor a model in which RSP4-containing head complexes are added to the preexisting spoke stalks of *pf1* flagella (Fig. 6A.

**Fig. 6).**
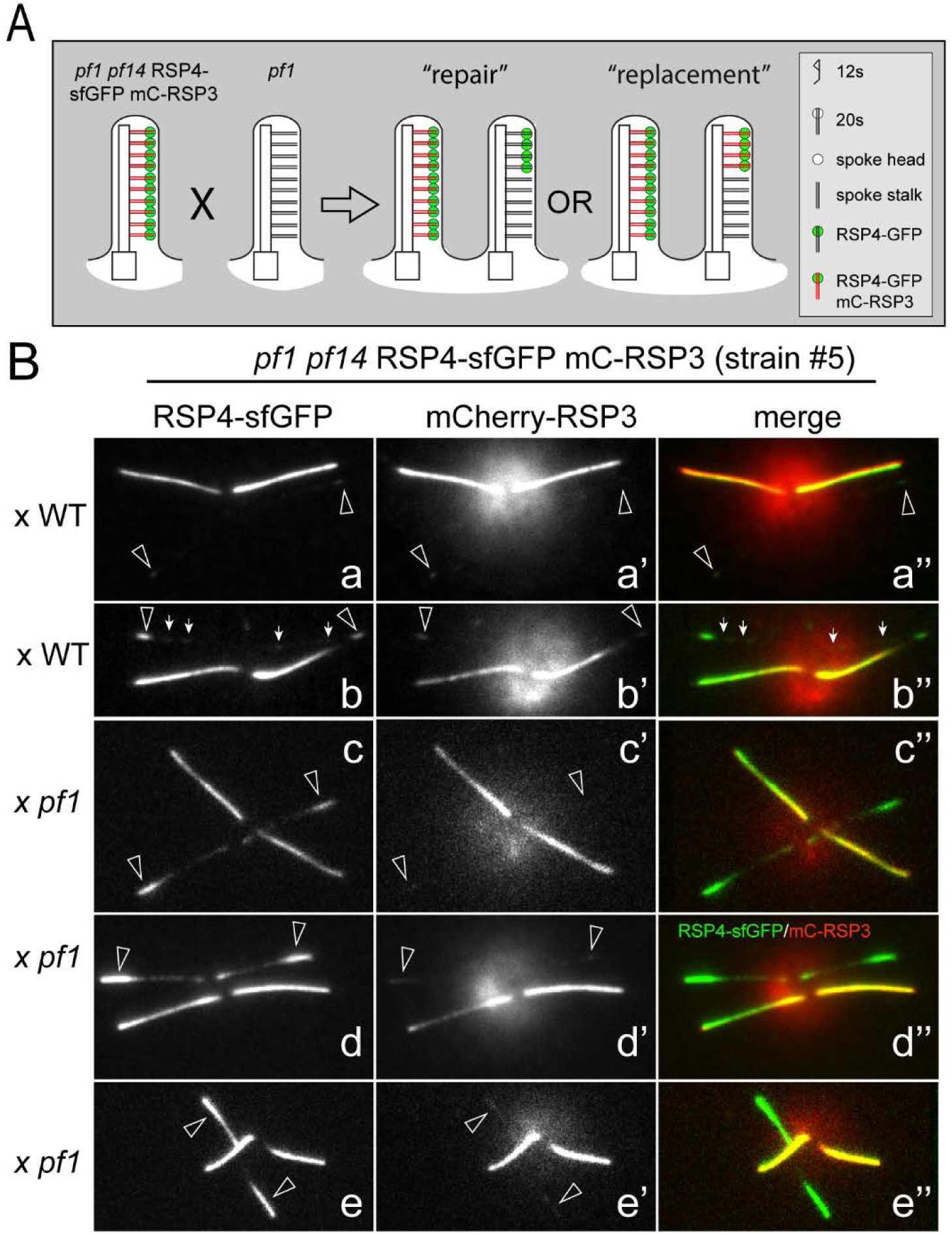
RSP4 assembles into spoke head-deficient pf14 flagella independently of RSP3. A) Schematic presentation of the mating between the double strain (*pf1 pf14* RSP4-sfGFP mC-RSP3) and *pf1*, and the possible outcomes, spoke repair by addition of RSP4-sfGFP heads or replacement with newly imported entire mC-RSP3 RSP4-sfGFP spokes. B) Gallery of images of zygotes obtained by mating the double mutant double rescue strain to either wild-type (WT) or *pf1*. Shown are the sfGFP (a – e), mC (a’ – e’), and merged (a” – e”) images of life cells. Arrowheads indicate the acceptor flagella. Small arrows, RSP4-sfGFP present in wild type-derived flagella. The incorporation of mC-RSP3 is generally limited to the flagellar tip. Note near absence of mC-RSP3 from the pf14-derived flagella of late zygotes, which possess RSP4-sfGFP along their entire length (e). Bar = 2 μm.

### IFT of RSP4-sfGFP requires RSP3

The observation that RSP4-sfGFP incorporates into *pf1* flagella without the apparent exchange of untagged RSP3 with mC-RSP3 could be explained by the transport of a RSP4-sfGFP complex independently of RSP3. First, we wanted to ensure that RSP3 and RSP4, which are both present in the 12S RS precursor complex, are indeed co-transported. RSP3 and RSP4 are thought to be dimers in the mature spokes and probably the 12S precursor (Wirschell et al., 2008). While mNG and sfGFP tagged version of RSP3 and RSP4 were readily detectable during transport, we were not able to detect mC-RSP3transport, which could be explained by the low brightness and stability of mC. During the course of this study, mScarlet I (mS), a novel bright monomeric red fluorescent protein, became available (Bindels et al., 2017). Using mating, we generated a strain expressing mS-RSP3 and mNG-RSP4 in the corresponding double mutant background (Fig. 7A). While still challenging, co-transport of tagged RSP3 and RSP4 was observed in regenerating cilia (Fig. 7B, C). In 48 out of 60 analyzed IFT events, mS-RSP3 co-migrated with RSP4-sfGFP including seven retrograde transports, six mS-RSP3 transports lacked apparent RSP4-sfGFP, and for six mS-RSP3 transports it was unclear whether mNG-RSP4 was associated or not. Co-migration of RSP3 and RSP4 was also observed in longer flagella, when only a small share of the IFT trains carries RS proteins. The data support the notion that RSP3 is mostly in a complex with RSP4 during IFT transport. However, we noted numerous mNG-RSP4 transports without detectable co-migrating mS-RSP3 signal. The lack of mS-RSP3 signals is likely caused by technical reasons, i.e., the comparatively low brightness and photostability of mS compared to mNG; also mNG folds in <10 minutes while mS requires several hours to mature, which will prevent light emission from mS-RSP3 newly translated during flagellar regeneration (Bindels et al., 2017; Shaner et al., 2013).

**Figure 7).**
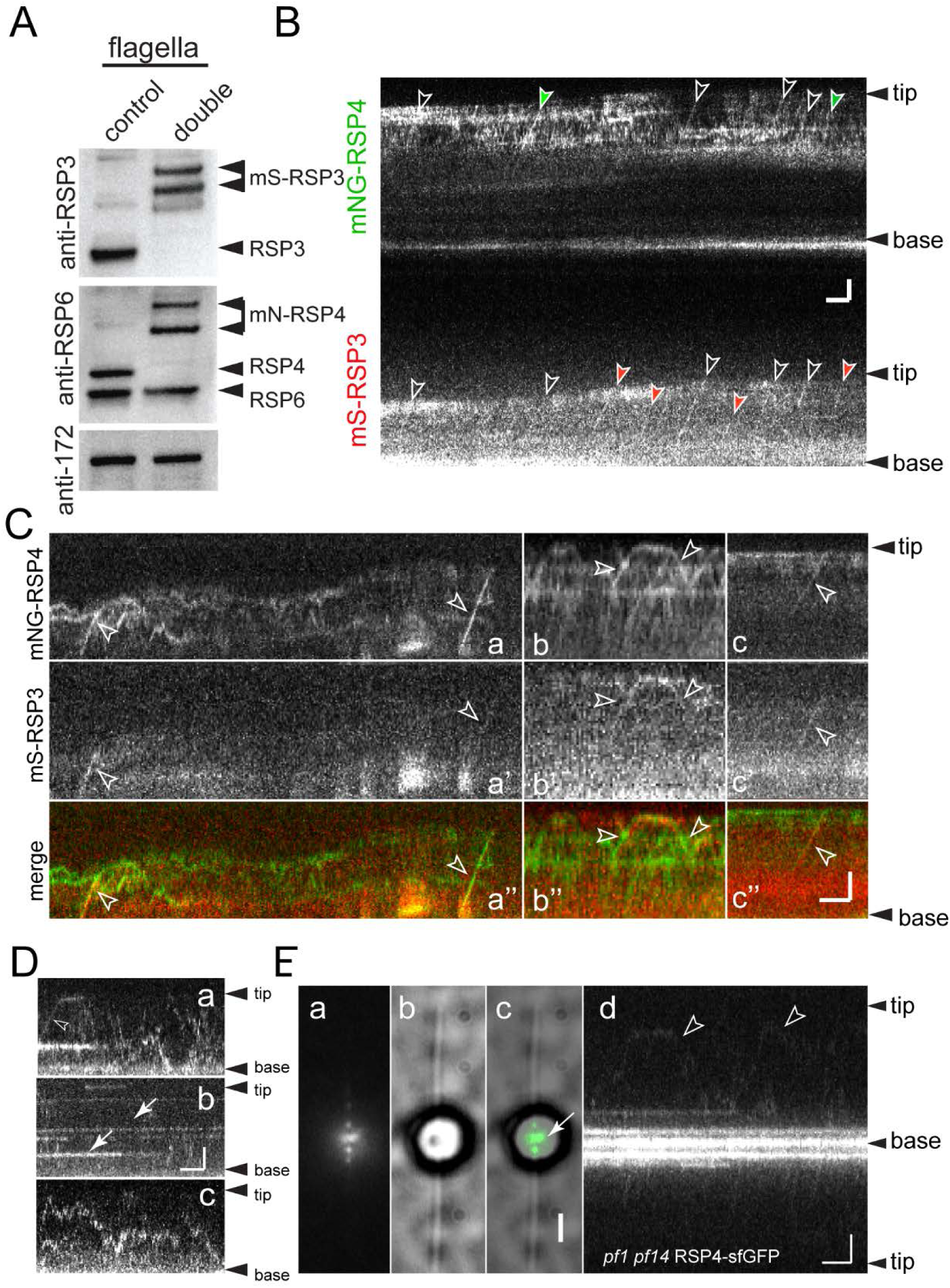
In the absence o/RSP3, RSP4-sfGFP accumulates at the flagellar base. A) Western blots analyzing flagella isolated from a control strain of the *pf1 pf14* mS-RSP3 mNG-RSP4 double mutant double rescue strain. Two replica membranes were stained with anti-RSP3 and anti-RSP6 as indicated; anti-IC2 staining was used as a loading control. B) Raw kymogram from a simultaneous recording of mS-RSP3 and mNG-RSP4 in regenerating flagella. Co-transports are marked by open arrowheads. Colored arrowheads indicate transports visible only in one of the two channels. Bar = 2s 2 μm. C) Gallery of kymograms showing co-transport of mNG-RSP4 and mS-RSP3 in regenerating flagella. Bar = 2s 2 μm. D) Gallery of kymograms showing RSP4-sfGFP expressed in the *pf1 pf4* double mutant background in regenerating (a, c – d) and steady-state (b) flagella. Arrowhead, Anterograde IFT. Arrows, Stationary signals. Bar = 2s 2μm. E) TIRF (10-frame image average), bright field, and merged image of a live *pf1 pf14* cell showing the accumulation of RSP4-sfGFP near the basal bodies and in a spotted fashion in the proximal portions of the flagella. Arrowheads RSP4-sfGFP diffusion. Bars = 2s 2μm.

However, the observation could be also explained by RSP3-independent transport of RSP4. Indeed, Diener et al. (2011) reported the presence of a RSP4/6/9/10 complex in the cell body cytoplasm of *pf14* mutants indicating that near complete spoke heads assemble in the absence of RSP3. To test if such head complexes travel via IFT, we expressed RSP4-sfGFP in *pf1 pf14* double mutants (Fig. 7D, E). Anterograde and retrograde IFT of RSP4-sfGFP was sporadically observed in both steady state and regenerating flagella but its frequency was exceedingly low (<0.01 events/minutes; Fig. 7D a). RSP4-sfGFP occasionally remained stationary in *pf1 pf14* flagella indicative for association with the axoneme, either unspecifically or by transient interaction with the central pair (Fig. 7D b). Finally, RSP4-sfGFP often diffused inside the mutant flagella (Fig. 7D c, E). The data indicate that some RSP4-sfGFP can enter the flagella independently of RSP3. However, the incorporation of RSP4-sfGFP into *pf1*-derived mutant flagella occurs in a pronounced tip-to-base pattern presumably because the protein is shuttled via IFT to the tip, where it will be released and diffuse to nearby docking sites. We think that neither low frequency IFT nor entry of RSP4-sfGFP by diffusion is likely to explain that pattern observed during the repair of *pf1* flagella; RSP4-sfGFP entering *pf1* flagella by diffusion should bind first to vacant sites near the base of the cilium as described for *pf27*, a mutant lacking the putative RS-IFT adapter (Alford et al., 2013).

Interestingly, RSP4-sfGFP accumulated near the basal bodies of many *pf1 pf14* RSP4-sfGFP cells with steady state or regenerating flagella (Fig. 7E). In contrast to previous antibody-based studies, an accumulation of RSP4-sfGFP and RSP3-sfGFP was not observed in the corresponding *pf1* and *pf14-*derived rescue strains with growing or steady state flagella (not shown; (Qin et al., 2004)). Thus, the build-up of RSP4-sfGFP at the basal bodies is likely to be related to the near complete failure to associate to IFT in the absence of RSP3. In summary, in vivo imaging supports biochemical data that the 12s RS precursor complex encompassing RSP3 and RSP4 is transported by IFT.

## Discussion

Using direct *in vivo* imaging of FP-tagged RS proteins, we show that these proteins are transported by IFT into *Chlamydomonas* flagella. The observations confirm previous studies indicating transport of RS proteins to the flagellar tip by an active mechanism (Johnson and Rosenbaum, 1992). Also, RS proteins have been shown to co-IP with IFT proteins from flagellar extracts (Qin et al., 2004). Direct visualization of RS proteins during transport uncovered additional features of RS transport, which would be difficult to determine by biochemical approaches. Both RS proteins tested here showed a high propensity to move on IFT trains in one run from the flagellar base to the tip and the transport of both proteins was upregulated during flagellar elongation. The data are reminiscent of previous observations on the transport of FP-tagged tubulin and DRC4 and further support the notion that the amount of axonemal proteins present on an IFT train is upregulated during flagellar growth (Craft et al., 2015; Wren et al., 2013). The processivity of an IFT-cargo complex is likely to reflect the stability of the interaction. Because different cargoes attach to distinct sites of the IFT particle, we expect differences in the stability of the various IFT-cargo complexes (Bhogaraju et al., 2013; Lechtreck, 2015; Taschner et al., 2017). Indeed, different cargoes showed distinct profiles in the distribution of unloading sites along the flagellar length. Under the assumption that the distance traveled together reflects the stability of IFT-cargo complexes, RSP4-IFT is more stable than DRC4-IFT and less stable than tubulin-IFT.

Two-color life imaging has shown transport of various proteins on IFT trains. The low-number cargoes are typically tagged with bright and stable green-light emitting FPs whereas the IFT carriers were tagged with red fluorescing proteins such as mCherry, which are not only less bright and photostable but also more difficult to image due to the bright red autofluorescence of *C. reinhardtii*. These shortcomings are mitigated by the fact that IFT trains are repetitive structures consisting of 15 - 40 IFT particles corresponding to a similar number of red FPs. Here, we accomplished simultaneous imaging of RSP3 and RSP4, two low number cargo proteins. Even when transport of RSs occurred at a low frequency as it is the case in longer regenerating flagella, most mS-RSP3 traveled in association with mNG-RSP4 on the same IFT train. This suggests that both proteins are in a complex rather than being associated independently of each other to different subunits of the IFT train. In comparison to other axonemal proteins studied in our laboratory, both RS proteins were very difficult to image presumably indicating a low number of active FPs in the transport unit.

Our *in vivo* imaging data are in agreement with previous biochemical data demonstrating a 12S complex encompassing RSP3 and RSP4 in the cell body and flagellar matrix. Inside cilia, the 12S RS precursor needs be transformed into the mature RSs; during this process, additional RS proteins are recruited into the complex but most steps of the conversion mechanism remain unclear. It has been suggested that two of the Γ-shaped 12S complexes will dimerize to form the scaffold of the largely symmetric T-shaped RSs (Pigino and Ishikawa, 2012). Alternatively, the 12S precursor could be remodeled into the 20S complex, e.g., by recruiting HSP40 and the dimeric 10-kDa protein LC8; LC8, for example, could bind and stretch-out RSP3 dimers already present in the 12S precursor (Gupta et al., 2012).

The dimensions of the 12S complex (~28nm length and 20nm width) and of 20S spokes isolated from axonemes (50nm length and 25nm width, each based on negative stain) support the notion that more complex rearrangements are required to convert the 12S into the 20S complex (Diener et al., 2011).

The current models of RS assembly assume that RSP3 and RSP4 present in a given 12S precursor complex are not only transported together but also remain together during the conversion into mature spokes. However, during the repair of the head-less stalks in *pf1*-derived flagella of dikaryons, RSP4-sfGFP provided by the *pf1 pf14* RSP4-sfGFP mC-RSP3 donor strain incorporated without a matching incorporation of mC-RSP3. This pattern suggests that RSP4-sfGFP separates at one point from mC-RSP3 and is then added onto the existing untagged spoke stalks of the *pf1* axoneme. We also observed that the incorporation of RSP4-sfGFP along the length of wild-type derived zygotic cilia exceeded that of mC-RSP3. This indicates that untagged spoke heads were exchanged with newly imported tagged ones while the untagged RS stalk remained in place.

One caveat of our approach is that the mC and mS tags are considerably weaker and less stable than the sfGFP and mNG tags; thus, weaker FP-RSP3 signals could have been missed detection. However, late zygotes possessed plenty of newly assembled RSP4-sfGFP along their entire length of the *pf1*-derived acceptor flagella and both FP-proteins were prominently visible in the two donor flagella and often near the tip of the acceptor flagella. Thus, the photochemical properties of the FPs are unlikely to explain the near absence of mC-RSP3 signals in post-repair *pf1*-derived acceptor flagella. The second caveat arises from the presence of untagged endogenous RSP3 in the zygotes. Cells could favor untagged over tagged RSP3 during the assembly or transport of RS precursors. However, dual tagged RSP3-RSP4 complexes were present and largely functional in the donor cell flagella. Further, both RSP4-sfGFP and mC-RSP3 were observed near the tip of zygotic acceptor flagella demonstrating transport and incorporation of mC-RSP3 even in the presence of endogenous RSP3 and our ability to detect such signals.

It is unclear whether the proposed split of the 12S complex occurs only during the repair of head-less *pf1* spokes and slow turnover of spoke heads in flagella with intact RSs or also during RS *de novo* assembly in growing cilia and the repair of *pf14* flagella lacking spokes. During the repair of spoke-less *pf14* flagella, tagged RSP3 and RSP4 appeared over the entire length of cilia with a somewhat stronger signal near the tip. This observation is distinct from the tip-to-base repair pattern previously described for RSP3 and RSP2 in *pf14* flagella based on antibody staining. We explain the differences with the increased sensitivity of *in vivo* TIRF of unfixed cilia while immunofluorescence could accentuate the somewhat higher density of RS proteins near the tip. In contrast, RSP4-FP is assembled in a pronounced tip-to-base pattern onto *pf1* axonemes. However, both FP-tagged RS proteins mostly dissociate from IFT at the ciliary tip from where they move into the ciliary shaft by diffusion. Thus, the distinct pattern described above could indicate that the RSP4-containing head complexes are assembly-competent once they leave the tip region and will diffuse and dock to the next available head-less stalk they encounter. RSP3-complexes, however, need more time to mature and thus will diffuse deeper down into the cilium before docking.

In summary, our data suggest that RSP3 and RSP4 enter cilia in a precomplex but separate into a RSP3- and RSP4-containing subcomplexes before the formation of mature spokes. Co-transport of RS proteins could ensure that they enter cilia in the proper stoichiometry. Also, IFT particles possess a limited number of cargo binding sites favoring transport of complexes in which only one or a few of the subunits directly interact with IFT. Within cilia, the separation of RSP3- and RSP4-subcomplexes could aid in axonemal maintenance by facilitating the replacement of damaged or lost spoke heads. Current data revealed similarities and differences in assembly, transport, and final mounting of axonemal dynein and RSs; the future analysis of additional axonemal substructures promises to reveal additional processes utilized to assemble axonemal substructures.

## Acknowledgements

This study was supported by the National Institutes of Health (R01GM110413 to K.L.). The content is solely the responsibility of the authors and does not necessarily represent the official views of the National Institutes of Health. The authors declare that they have no conflict of interest.

## Materials and Methods

All strain were maintained in Minimal (M) medium (https://www.chlamycollection.org/methods/media-recipes/minimal-or-m-medium-and-derivatives-sager-granick/) at 24 C with a light/dark cycle of 14/10 hours. The following strains were obtained from the Chlamydomonas Resource Center (https://www.chlamycollection.org/): CC-613 (pf14 mt−), CC-1032 (pf14 mt+), CC-1024 (pf1 mt+), CC-602 (pf1 mt−), CC-620 (nit1 nit mt+), and CC-621 (nit1 nit2 mt−).

For C-terminal tagging of RSP4 with super folder GFP (GFP), *RSP4* was amplified by PCR using the primers RSP4f (CGCCTCGAGATGGCGGCAGTGGACAGC) and RSP4r (CGCAGATCTGCTGCCGCCGCCGCTG) and the genomic RSP4-sfGFP construct described by ODA et al. 2011 as a template. The cleaned PCR product was digested with Xho1 and BlgII, gel-purified, and ligated into the pBR25-sfGFP-sfGFP vector (Lechtreck, 2016) digested with Xho1 and BamH1. This placed the RSP4 CDS upstream of the gene encoding GFP. For N-terminal tagging of RSP4, the coding sequence was amplified using the primers RSP4-f-Bg12 (CGCAGATCTGGCGGCAGCGGCGGCATGGCGGCAGTGGACAGCG) and RSP4-r-EcoR1 (GCGGAATTCCTACTCGTCCGCCTCGGCCTC) and ligated downstream of the mNeonGreen (mNG) gene using the pBR25-mNG-α-tubulin vector described in CRAFT et al. For N-terminal tagging of RSP3, the *RSP3* coding region was amplified from a RSP3 cDNA clone using the primers RSP3f (CGCGGATCCATGGTGCAGGCTAAGGCGCA) and RSP3r (GCGGAATTCTTACGCGCCCTCCGCCTC). After digestion with BamH1 and EcoR1, the PCR fragment was gel-purified and ligated into the pBR25-mNeon-α-tubulin vector digested with the same enzymes (Craft et al., 2015). Derivates were obtained by replacing the mNeon gene with genes encoding other FPs by restriction digest and ligation. Primers mScarlet-f-XhoI (GCGCTCGAGATGGTGAGCAAGGGCGAG) and mScarlet-r-BamHI (CGCGGATCCCTTGTACAGCTCGTCCATGCC) were used to amplify mScarlet-I (mS) using a template codon-adopted for expression in mammalian cells. For transformation, the plasmids were digested with Xba1 and Kpn1, the ble-RSP4-sfGFP and ble-mNeon-RSP3 cassettes were gel purified, and introduced into appropriate strains (CC-602 and CC-1032) by electroporation. Transformants were selected on zeocin plates (10 μg/ml), transferred to liquid medium, and analyzed for restoration of motility. Expression of FP-tagged RS proteins was verified by TIRF microscopy and Western blotting. A strain expressing both RSP4-sfGFP and mCherry (mC)-RSP3 was obtained by mating the corresponding single rescue strains in CC-602 (RSP4-sfGFP *pf1* mt −) and CC-1032 (mC-RSP3 *pf14* mt+). The progeny was screened by TRIF microcopy for expression of GFP and mC and Western blotting was used to obtain *pf1 pf14* RSP4-sfGFP mC-RSP3 double mutant double rescue strain. The *pf1 pf14* RSP4-sfGFP strain was isolated from the progeny of this cross. The *pf1 pf14* mNG-RSP4 mScarlet-RSP3 double mutant double rescue strain was generated following the same strategy.

For Western blot analyses, cells in HSM were deflagellated by the addition of dibucaine. After removing the cell bodies by two differential centrifugations, flagella were sedimented 40,000 x g, 20 minutes, 4 C°. Flagella were dissolved in Laemmli SDS sample buffer, separated on Mini-Protean TGX gradient gels (BioRad), and transferred electrophoretically to PVDF membrane. After blocking, the membranes were incubated overnight in the primary antibodies; secondary antibodies were applied for 90–120 minutes at room temperature. After addition of the substrate (Femtoglow; Michigan Diagnostics), chemiluminescent signals were documented using A BioRad Chemi Doc imaging system. The following primary antibodies were used in this study: rabbit anti-RSP6 (Williams et al., 1986), rabbit anti-RSP3 (Williams et al., 1989), rabbit anti-GFP (A-11122; ThermoFischer), mouse monoclonal anti-IFT172 (Cole et al., 1998), and mouse monoclonal anti-IC2 (King and Witman, 1990).

Live imaging was performed on a Nikon eclipse Ti-U equipped with a 60x NA1.49 TIRF objective, 488-nm and 561-nm diode lasers (Spectraphysics), a dual view system (Photometrics), and an EMCCD camera (Andor iXon3). Data were collected using the Nikon Elements software package and exported into Fiji for further analysis including kymography and frame extraction. Contrast and brightness were adjusted in Fiji and in Photoshop and all figures were prepared in Illustrator (Adobe). Specimen preparation, TIRF imaging, and data analysis have been described in detail in (Lechtreck, 2013) and (Lechtreck, 2016).

For flagellar regeneration experiments, 0.5 M acetic acid was rapidly added to cells in fresh M medium under stirring (1,200 rpm) to reach a pH of ~4.25. After ~45s, KOH (0.25 M) was added to neutralize the solution (pH ~6.8), the cells were collected by centrifugation, resuspended in M medium and either used directly or stored on ice until needed. During regeneration, cells were incubated on a shaker in bright light at room temperature. Specimens for live imaging were prepared 25 minutes after the pH shock and later.

For mating experiments, cells were grown in bright light on M plates for ~9–12 days and then transferred to dim light for 2–3days (see above for conditions). The evening before the experiment, cells were resuspended in M-N medium (7–10 ml/plate) and incubated overnight in constant light with agitation; agitation was omitted for the RS mutants, which assembled cilia better without shaking. In the morning, cells were transferred to 1/5M-N supplemented with 10 mM Hepes pH 7 and incubated for an additional ~2–6 hours. Gametes of opposite mating types were mixed in a microcentrifuge tube and samples for imaging were taken over a period of ~5 minutes to several hours.

## Abbreviations

IFT: intraflagellar transport
FP: fluorescent protein
mC: mCherry, mNG, mNeonGreen
mS: mScarlet-I
RS: radial spoke
RSP: radial spoke protein
sfGFP: superfolder GFP

